# Detecting differential transcription factor activity from ATAC-seq data

**DOI:** 10.1101/315622

**Authors:** Ignacio J. Tripodi, Mary A. Allen, Robin D. Dowell

## Abstract

Transcription factors are managers of the cellular factory, and key components to many diseases. Many non-coding single nucleotide polymorphisms affect transcription factors, either by directly altering the protein or its functional activity at individual binding sites. Here we first briefly summarize high throughput approaches to studying transcription factor activity. We then demonstrate, using published chromatin accessibility data (specifically ATAC-seq), that the genome wide profile of TF recognition motifs relative to regions of open chromatin can determine the key transcription factor altered by a perturbation. Our method of determining which TF are altered by a perturbation is simple, quick to implement and can be used when biological samples are limited. In the future, we envision this method could be applied to determining which TFs show altered activity in response to a wide variety of drugs and diseases.

## 1. Introduction

Transcription factors (TFs) are the managers of the cellular factory, controlling everything from cellular identity to response to external stimuli[1]. Because of their central importance in interpreting the genome, millions of people are affected by mutations residing within TFs[2], causing a wide variety of symptoms (see Table 1). For example, over half of all cancers have a mutation in the TF TP53[3].

**Table 1.**
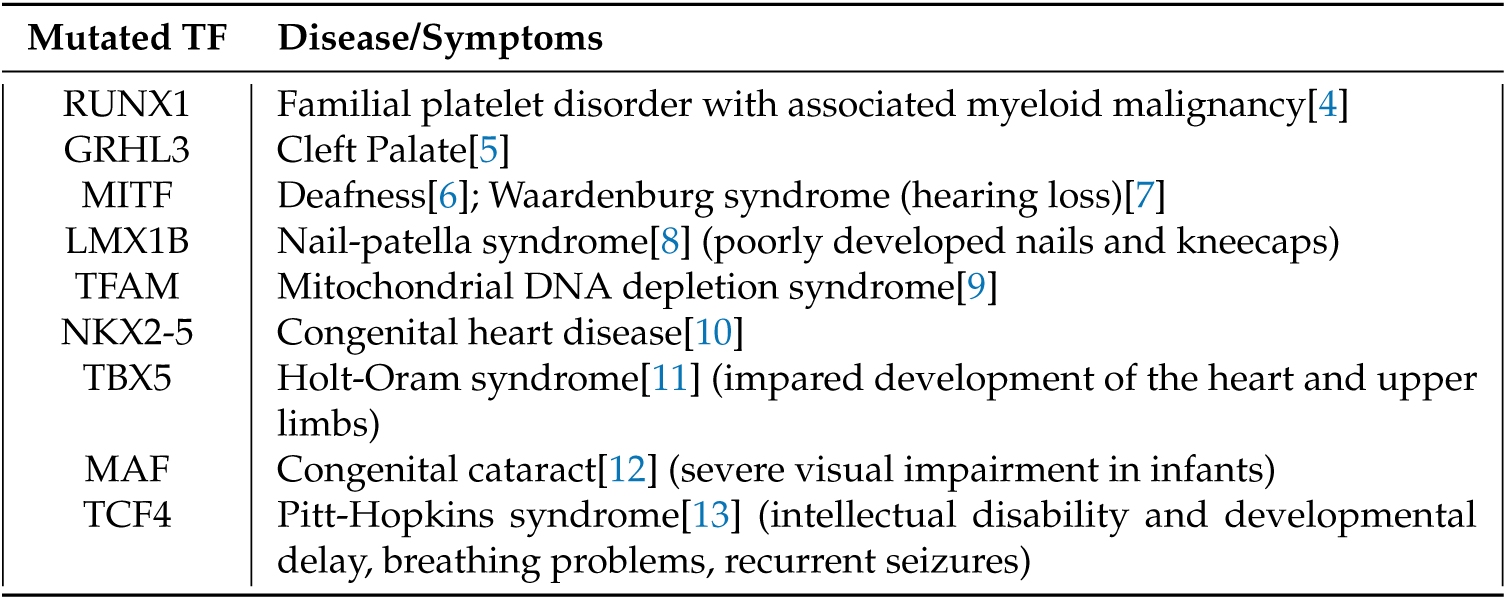
Examples of diseases caused by mutations in a transcription factor

Moreover, most disease-causing mutations are found in regulatory regions[14,15], e.g. enhancers, which are dense with TF binding sites[16]. A startling 60-76.5% of disease-associated single nucleotide polymorphisms (SNPs) are in enhancers[17–20], which are short regulatory regions densely bound by TFs[21]. In fact, the well known program HaploReg now lists all TFs that bind over each SNP, a useful piece of information for understanding the impact of a SNP[22].

The relationship between many diseases and transcription factors has led to tremendous interest in global investigations of transcription factor activity. To decipher transcription factor activity requires understanding the two major functions of a transcription factor: binding to DNA and modification of transcription. Transcription factors bind to specific DNA sequences, a TF recognition motif. A number of techniques have been utilized to identify and characterize these recognition motifs[23]. However, because most genomic instances of the motif are not actually bound, having the recognition motif is insufficient. Protein-DNA interactions can be measured genome wide using chromatin immunoprecipitation followed by sequencing (ChIP-seq)[23–25]. Unfortunately, numerous lines of evidence indicate that not all binding events influence transcription[26–28]. Conceptually, this is akin to saying that just because someone is standing in a lab (TF binding) it does not imply they are actually conducting an experiment (altering transcription). Therefore, distinct assays are necessary to identify the locations where a TF is bound to DNA and determine whether that DNA binding leads to altered transcription nearby. A number of high throughput assays are available to interrogate these two key functions.

Extensive attention has focused on determining where in the genome transcription factors bind[23, 29,30]. The ENCODE project alone included approximately 2000 TF ChIP-seq experiments, including 180 TFs in K562 (myeloid leukemia) cells alone[29]. Large regulation project such as ENCODE and Roadmap Epigenomics have been invaluable to our understanding of TF binding. However, there are an estimate 1600 TFs in the human genome and many do not have a reliable antibody for ChIP-seq[23]. Even when antibodies are available, individual transcription factors can have distinct profiles of binding locations across cell types and conditions. Consequently, the cost of individually profiling every TF in each cell type is enormous, much less across different conditions[31]. Finally, if the effect of a particular perturbation is unknown, profiling assorted TFs by ChIP is prohibitively expensive.

An alternative approach to detecting individual protein-DNA binding locations is to infer a large collection of binding events via DNA footprinting[32–34]. Dense mapping of DNase I clevage sites identifies small regions protected from cleavage by the presence of a bound transcription factor[32,33]. While early footprinting studies identified a large repertoire of previously un-characterized motifs protected from cleavage, suggesting many novel transcription factors[34], subsequent work indicates these regions likely reflect sequence based cleavage bias of the DNase I enzyme[35]. Additionally, it is also now clear that most TFs (80%) do not show a measurable footprint[36], thereby limiting the effectiveness of this approach.

Despite these limitations, DNA footprinting assays uncovered a distinct function for transcription factors: altering DNA accessibility. When chromatin accessibility data is considered in the context of known TF sequence motifs[37–40], one can reasonably infer transcription factor binding profiles[41,42]. When accessibility profiles are then compared to ChIP in the context of perturbations, transcription factors could be classified as “pioneer” or “settler” depending on whether they open chromatin or require accessible, exposed DNA to bind[42]. Whether alterations of local chromatin accessibility reflect a byproduct of the TF’s DNA binding or its altering of transcription remains unclear.

Altering transcription is the second major function of transcription factors[23]. Because TFs alter transcription, some of the earliest studies of TFs as regulators were based on expression data. For nearly twenty years large compendiums of expression data have been utilized to infer gene regulatory networks[43,44]. Typically these approaches search for modules, collections of co-regulated genes across distinct conditions. Identification of nearby TF recognition motifs[45,46] or co-regulated transcription factors[43] link particular TFs to the module of genes they regulate. For instance, ISMARA (Integrated System for Motif Activity Response Analysis)[47] models gene expression in terms of TF sequence motifs within proximal promoters. Gene regulatory network methods have been instrumental for understanding large scale regulatory networks, but are inherently limited by the fact that they depend on steady state expression data. Steady state expression assays (microarray or RNA-seq) reflect not only transcription but also RNA processing, maturation and stability. Hence, they are an indirect readout on the effect of perturbations to transcription factors. Additionally, they are generally incapable of reliably detecting small changes at short time points without an impractical number of replicates[48].

Nascent transcription assays (GRO-seq, PRO-seq) directly profile RNA associated with engaged cellular polymerases[49,50]. Consequently, nascent assays are a direct readout on changes to transcription induced by perturbations[21,51]. Interestingly, an additional feature of nascent transcription data is the identification of short unstable transcripts immediately proximal to sites of transcription factor binding[52–57]. Importantly, these transcripts, now known as eRNAs can be employed as markers of TF activity[58]. The change in patterns of eRNA usage genome-wide relative to TF recognition motifs allows one to determine which transcription factors are altered by a perturbation with no a priori information. Unfortunately, nascent transcription protocols[49,50] are onerous, time consuming, and require large numbers of cells as input. Consequently, these experimental assays are predominantly used on cultured cell lines and not yet widely adopted. Therefore, we sought a simpler, easy to use approach to inferring differential transcription factor activity.

The Assay for Transposase-Accessible Chromatin followed by sequencing (ATAC-seq), a method for identifying regions of open chromatin, is particularly attractive because it is quick, easy, inexpensive, and deployable in small cell count samples. Additionally, recent work has shown that changes in chromatin accessibility can inform on TF activity. Specifically, BagFoot[36] combined footprinting with differential accessibility to identify TFs associated with altered chromatin accessibility profiles in the presence of a perturbation. They predominantly focused on DNase I hypersensitivity data, but also examined a small number of ATAC-seq datasets. Here we seek to confirm and extend their results in two ways. First, we ask whether an alternative approach – namely the motif displacement statistic[58], developed initially for nascent transcription analysis, could infer differential TF activity from ATAC-seq datasets. Second, we sought to construct an easy-to-use pipeline specific to the analysis of differential ATAC-seq analysis.

## 2. Results

We introduce a tool, Differential ATAC-seq toolkit (DAStk), developed with simplicity and ease of implementation in mind, focused around inferring changes in TF activity from ATAC-seq data. Using nascent transcription data we had previously developed the motif displacement score (MD-score) a metric that assesses TF associated transcriptional activity. As such, the MD-score reflects the enrichment of TF sequence motif within an small radius (150 bp) of enhancer RNA (eRNA) origins relative to a larger local window (1500 bp)[58]. While ATAC-seq does not directly provide information on eRNAs, most sites of eRNA activity reside within open chromatin[59]. Therefore we utilize the midpoint of called ATAC-seq peaks (rather than the eRNA origin) as a frame of reference for calculating MD-scores. Then, given two distinct biological conditions, we compare the ratio of MD-scores across the conditions and identify statistically significant changes by a two-proportion Z-test. Using public ATAC-seq data from a variety of human and mouse cell lines (IMR90, H524, NJH29, ZHBTC4) and perturbations (nutlin, doxycycline, tamoxifen), we assessed changes in accessibility over all putative TF sequence recognition motifs (for all motifs within the HOCOMOCO database[38]).

Given our familiarity with TP53 activation[55,60], we first examined this approach on ATAC-seq data gathered before and 6 hours after Nutlin-3a exposure on IMR90 cells[61]. Nutlin-3a is an exquisitely specific activator of TP53. As expected, we found that TP53 displayed the most significant change (p-value < 10 −5) in MD-score (Figure 1A, in red) of all motifs within the HOCOMOCO database[38]. Relaxing the p-value cutoff (p-value < 10−4) we subsequently identified altered activity in TP63 and TP73 (Figure 1A, in maroon), likely reflecting the fact that these two proteins have nearly identical sequence recognition motifs as TP53.

**Figure 1.**
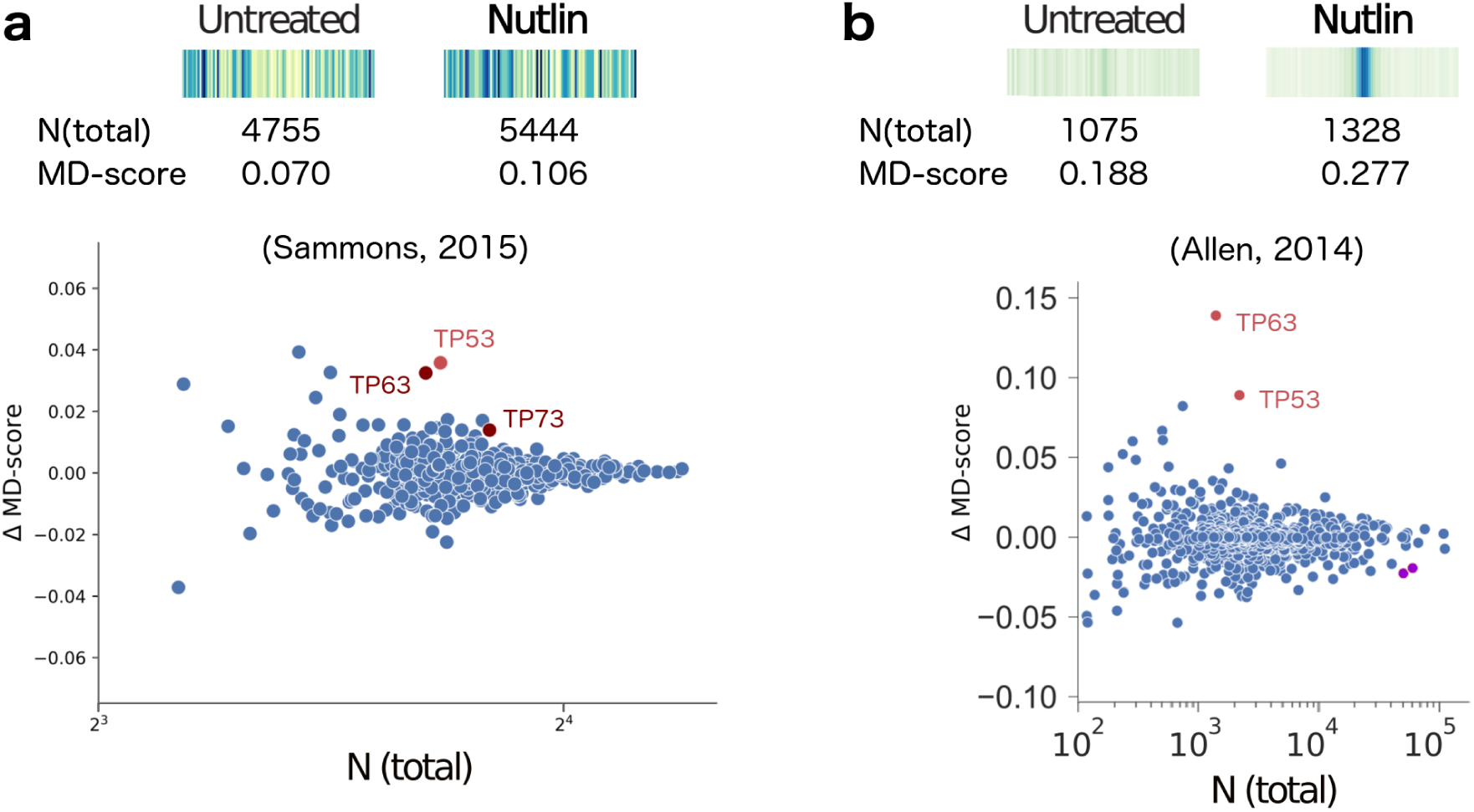
(**a**)Top: The motif displacement distribution as heatmap (increasingly dark blue indicates more instances of motif), MD-score and the number of motifs within 1.5kb of an ATAC-seq peak before and after stimulation with Nutlin-3a (e.g. Nutlin)[61] for TP53, the transcription factor known to be activated. Bottom: For all motif models (each dot), the change in MD-score following perturbation (y-axis) relative to the number of motifs within 1.5kb of any ATAC-seq peak center (x-axis). Red/maroon points indicate significantly increased MD-scores (p-value < 10^−5^, < 10^−4^, respectively). (**b**) Similar analysis obtained from nascent transcription data[55], where MD-scores are measured relative to eRNA origins. Purple dots indicate significantly decreased MD-scores. Figure adapted from Azofeifa et al [58].

Interestingly, Nutlin-3a has also been analyzed using nascent transcription data albeit in a different cell line (HCT116) at a shorter time point (1 hour)[55]. The MD-score analysis of the nascent data[58] obtained very similar results (Figure 1B). Unfortunately, a direct comparison of individual genomic loci between the two data sets is not feasible because they used different cell lines and drug exposure times. However, a couple of interesting observations concerning the overall MD-score trends are none-the-less noteworthy. First, the co-localization of the TP53 motif with ATAC-peak midpoints is far less striking than the co-localization of motifs with the eRNA origins (observed in the heatmap histograms). This observation, combined with the relative lower magnitude of ∆MD-scores (y-axis) suggests that the eRNA origin (obtained in nascent transcription) is a far more precise method of localizing and detecting changes in TF activity. Second, despite this lack of precision, ATAC-seq correctly identifies TP53 as the most dramatically altered MD-score whereas the best scoring motif with nascent transcription is TP63. Why this discrepancy exists is unclear, but given the relative similarity of these two motifs it may simply be coincidental.

We next analyzed differential ATAC-data gathered by Denny et. al to examine whether Nfib promotes metastasis via increasing chromatin accessibility. For this question, they examined two human small cell lung carcinoma (SCLC) cell lines (H524 and NJH29), profiling by ATAC-seq before and four hours after doxycycline treatment. Using the MD-score approach, we detect changes in TF activity for multiple members of the NFI family (Figure 2A,B). An increase in NFIA (two different motifs) and NFIC was detected in both cell types (p-value < 10^−5^ for H524s; p-value < 10^−10^ for NJH29s). As further confirmation of the NFI signal, we tested one of their mouse samples (KP22 cells) and found an increase of NFIA (p-value < 10^−5^), consistent with the human results. We next asked whether our results were sensitive to the particular peaks utilized. To this end we sub-sampled peaks from the NJH29 data and re-ran our analysis. Both NFIA and NFIC are detectable as significant (p-value < 10^−10^) even when using only half of the ATAC-seq peaks, suggesting the signal is reasonably robust.

**Figure 2.**
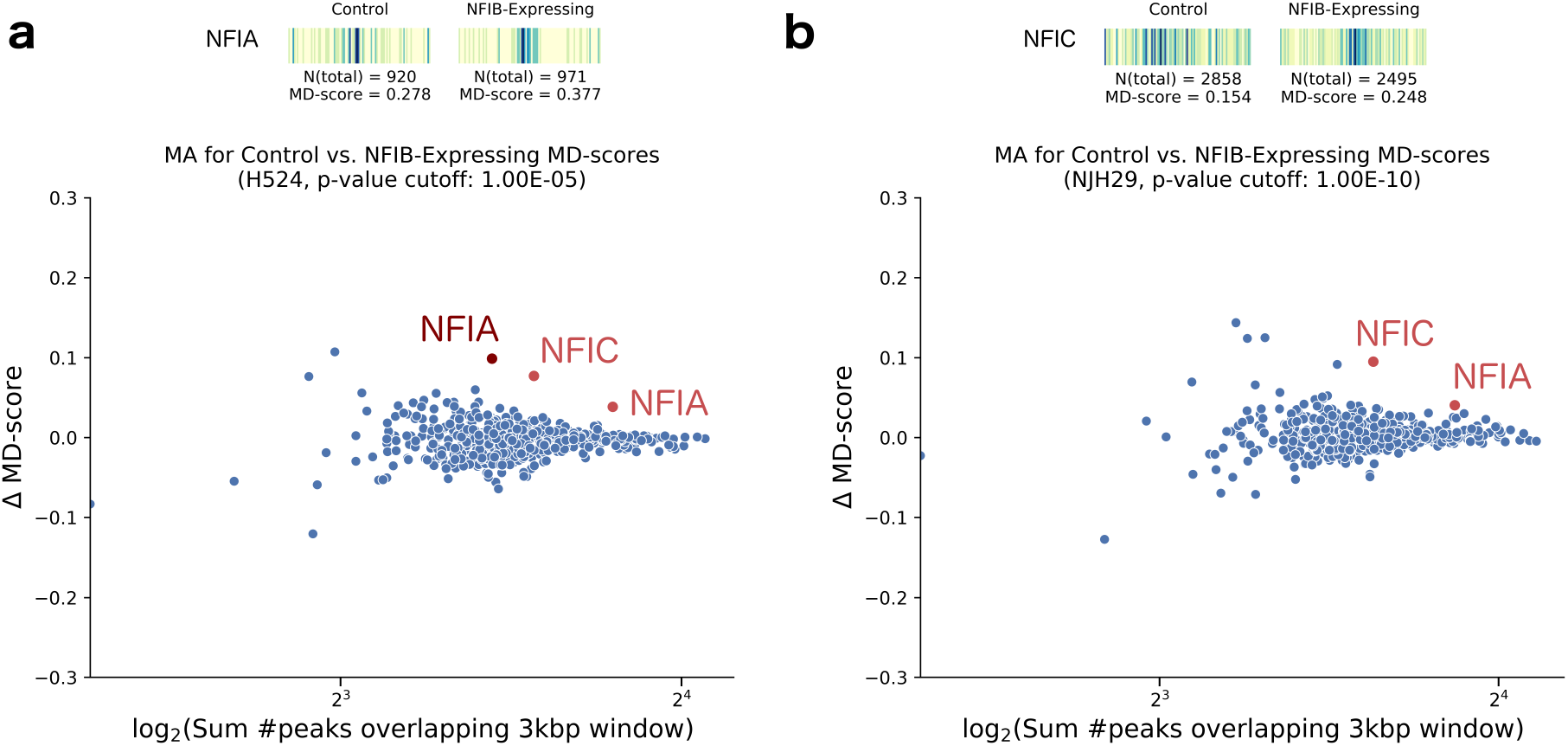
(**a**) Top: Motif displacement distribution as heatmap (increasingly dark blue indicates more instances of motif), MD-score and the number of motifs within 1.5kb of an ATAC-seq peak in control and NFIB-induced H524 cells with doxycycline, for the upregulated TF NFIA. Bottom: For all motif models (each dot), the change in MD-score following perturbation (y-axis) relative to the number of motifs within 1.5kb of any ATAC-seq peak center (x-axis). Red/maroon points indicate significantly increased MD-scores (p-value < 10-5, < 10-4, respectively). (**b**) Equivalent analysis performed on NJH29 cells, displaying a motif displacement distribution of the NFIC TF upregulation. We note that in the doxycycline-treated cells, most ATAC-seq peaks are located closer to the motif center than on the control cells.

We then sought to determine how the ∆MD-score approach compared to the BagFoot[36] at identifying differential TF activity. BagFoot also identified NIFA and NIFC within the SCLC differential ATAC-seq data[36]. However, they additionally claimed HNF6 as potentially altered in the SCLC data. Importantly, Baek et. al. noted that the HNF6 result did not hold when their approach utilized bias corrected data (based on naked DNA digested with Tn5). Given our MD-score approach does not identify HNF6 as altered further supports the idea that this result reflects a data artifact rather than a true biological phenomena. Interestingly, the MD-score approach and Bagfoot obtained nearly identical results on a second differential ATAC-seq dataset. In this case, King and Klose[62] showed BRG1, essential for pluripotency-related chromatin modifications, is required to make chromatin accessible at OCT4 target sites. To this end they treated ZHBTC4 mouse embryonic stem cells (ESCs) with tamoxifen for 72 hours to block BRG1 expression. When compared to the unperturbed mouse ESC control, we observed lowered MD-scores for SOX2, PO5F1 (Oct4) and NANOG in the BRG1-blocked cells (p-value < 10^−13^; Figure 3A), directly confirming the BagFoot findings.

**Figure 3.**
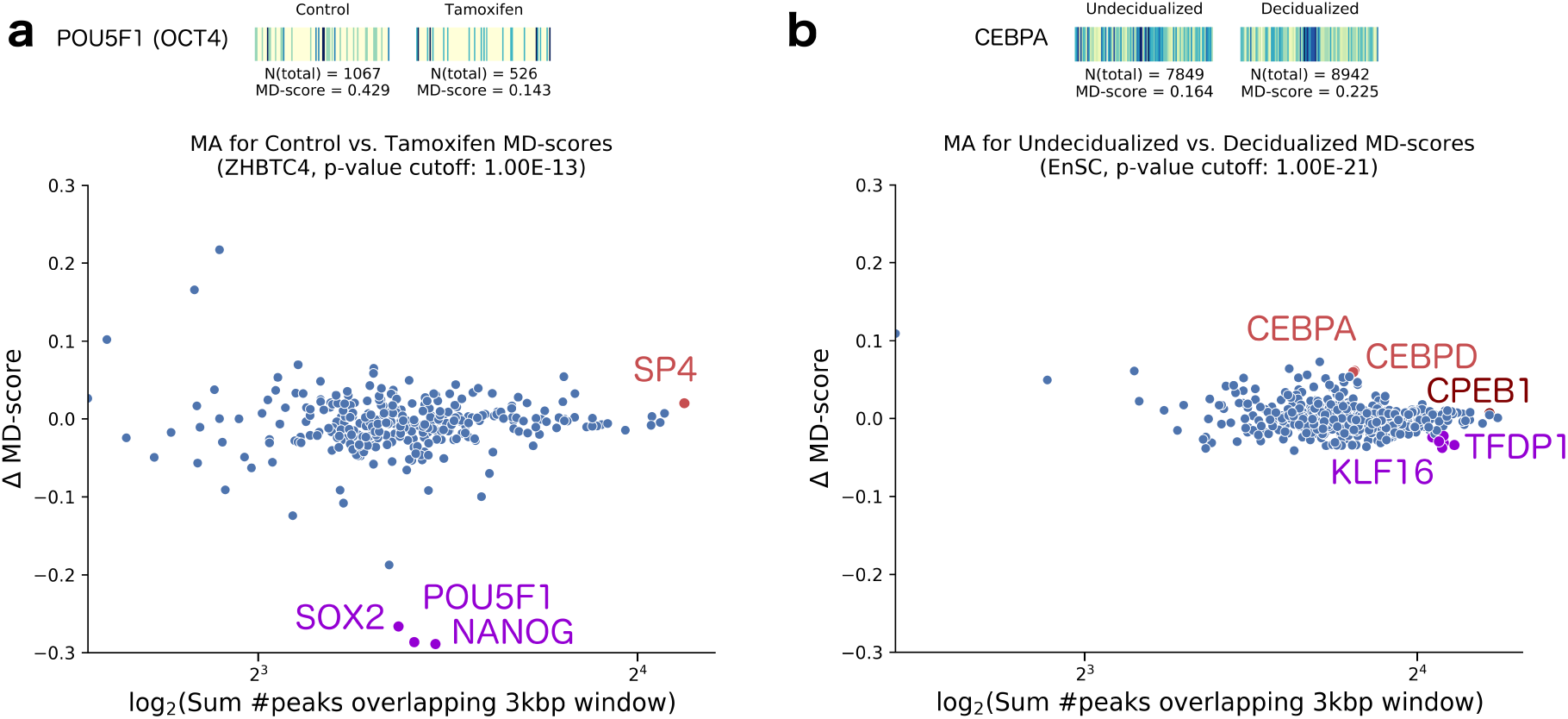
(**a**) Top: Motif displacement distribution as heatmap (increasingly dark blue indicates more instances of motif), MD-score and the number of motifs within 1.5kb of an ATAC-seq peak before and after stimulation with Tamoxifen[62] for the inhibited TF POU5F1, also known as OCT4. We observe that the decreased MD-score reflects not only a smaller number of peaks nearby this motif, but also a sharp decrease in co-localization with the motif. Bottom: For all motif models (each dot), the change in MD-score following perturbation (y-axis) relative to the number of motifs within 1.5kb of any ATAC-seq peak center (x-axis). Red points indicate significantly increased MD-scores (p-value < 10^−13^). Purple dots indicate significantly decreased MD-scores, at the same indicated p-value. (**b**) Equivalent analysis performed on endometrial stromal cells, before and after undergoing a decidualization process[63]. The motif displacement heatmap illustrates ATAC-seq peak distances to CEBPA, the TF expected to be upregulated.

Finally, we also examined a differential ATAC-seq data obtained for decidualized and undecidualized human endometrium cells[63]. Spontaneous decidualization occurs in response to progesterone signalling (i.e. by an implanted embryo at the early stages of pregnancy). Using our MD-score approach, we found the CEBP family of transcription factors had increased activity in decidualized cells, consistent with the author’s conclusion (Figure 3B). Additionally, we also found significantly lowered MD-scores for the KLF16 motif (a TF known to be involved in regulatory uterine cell biology[64]) and TFDP1 (a known target to the estrogen receptor *ERβ* present in all endometrial cell types[65] of lower activity during the secretory phase, in concert with the decidualization process). In all cases, the magnitude of MD-score alterations were relatively small, and yet the transcription factors uncovered can be linked to the underlying decidualization process.

## 3. Discussion

We sought to identify changes in TF activity across differential ATAC-seq datasets, as this protocol is inexpensive, simple and requires relatively small cell counts. Here we demonstrate two important results. First, using a simple statistic (the motif displacement score) as a co-localization measure of ATAC-seq peak midpoints to TF sequence motif sites across the genome, we correctly detect changes in TF activity. Second, our approach independently confirms the results obtained by BagFoot[36], as the two analysis techniques are distinct in their approach to quantifying differences in chromatin accessibility across conditions. Arguably, regardless of which analysis technique is preferred — differential ATAC-seq is a relatively simple and inexpensive way to assess for changes in TF activity induced by perturbations.

We believe there are two distinct advantages to the MD-score approach to assessing TF activity. First, the MD-score is calculated relative to a local background window. Consequently it cleanly accounts for the localized sequence bias observed at promoters and enhancers[58], which likely reduces false positives. Second, the statistic is relatively simple to implement and naturally accommodates multiprocessing for faster computations. DAStk can easily be incorporated at the tail-end of a traditional processing pipeline for ATAC-seq data, in that MD-scores are calculated directly from called peaks and genomic sequence.

Our MD-score statistic was originally developed for analysis of nascent transcription data[58] and focused on enhancer RNA co-localization with motifs. Given most eRNAs originate from areas of open chromatin[21,57,66] and many transcription factors can alter chromatin accessibility[42], it is perhaps unsurprising that differential chromatin accessibility can be used to infer changes in TF activity. However, it remains unclear whether the observed alterations of chromatin reflect a distinct functional activity of transcription factors or are simply a side effect of DNA binding and/or altering transcription. While a careful examination of the two Nutlin-3a datasets (Figure 1) identifies several genomic regions altered uniquely in only one of the two datasets (ATAC-seq or nascent), the lack of matched data makes interpretation of these differences difficult. Do they reflect differences of cell type or distinct functional activities of TP53? A careful comparison of chromatin accessibility and nascent transcription data in the context of a perturbation will be necessary to fully address this question.

## 4. Materials and Methods

### 4.1. Processing pipeline

Each ATAC-seq dataset was subjected to a standard data processing pipeline. The SRR datasets were converted to FASTQ format using `fastq-dump` v2.8.0 with argument –`split-3`. Paired-ended raw reads were trimmed using `trimmomatic` v0.36 at a fixed length with options PE -phred33 `CROP:36 HEADCROP:6`. After verifying the dataset quality with `FastQC` v0.11.5, the reads were aligned to the hg19 or mm10 reference genome, using `Bowtie` v2.2.9 with arguments `-p32 -X2000`. The resulting SAM files were converted to BAM format using `samtools` v1.3.1 using the `view -q` 20 -S -b arguments and sorted with the sort `-m500G` arguments. Bam files were then converted to BedGraph format for easier processing using bedtools v2.25.0 with arguments `-bg -ibam INPUT_BAM_FILE` -g `GENOME_REFERENCE` and read counts were normalized by the millions mapped. Finally, MACS v2.1.1.20160309 was used to call broad peaks from the ATAC-seq BAM files with arguments callpeak -n `ASSAY_PREFIX` -nomodel -format `BAMPE` -shift -100 -`extsize 200 -B -broad`.

The human motif sites calculated in Azofeifa et al[58] for the hg19 reference genome were used for human cells. The motif sites for mouse cells were obtained using FIMO with position weight matrices (PWMs) from HOCOMOCO, with a p-value cutoff of 10^−6^ (arguments -`max-stored-scores 10000000 –thresh 1e-6`.

### 4.2. Public Datasets

We used samples from the following public GEO datasets for our analysis: GSE58740 (samples SRR1448793 and SRR1448795), GSE81255/GSE81258 (samples SRR3493647, SRR3493653, SRR3493643, SRR3493645, SRR3493626, SRR3493627, SRR3493634, and SRR3493635), GSE87822 (samples SRR4413799 and SRR4413811), and GSE104720 (samples SRR6148318 and SRR6148319).

### 4.3. DAStk Software

The Differential ATAC-seq toolkit (DAStk) is a collection of scripts publicly available at https://biof-git.colorado.edu/dowelllab/DAStk for download. We used 642 PWMs of human motifs in the HOCOMOCO[38] database (to verify the presence of ATAC-seq peaks nearby), and 427 mouse motifs. TF sequence motifs were mapped to the hg19 or mm10 reference genomes with a p-value cutoff of 10^−6^. For each motif, the number of ATAC-seq peaks was accounted for, within a large (1500bp radius) and small (150bp radius) window, to calculate the motif displacement score. The difference between the MD-score in each condition and the number of ATAC-seq peaks nearby (large window) each motif were used to produce the MA plots. Those motifs with a statistically significant difference in MD-score were labeled, as determined by a z-test of two proportions[58].

## Acknowledgments

This work was funded in part by a NSF IGERT grant number 1144807 (IJT, RDD), NIH training grant T15LM009451 (IJT), a Sie Fellowship (MAA) and a NSF ABI DBI-12624L0 (RDD). The authors acknowledge the BioFrontiers Computing Core at the University of Colorado Boulder for providing High Performance Computing resources (NIH 1S10OD012300) supported by BioFrontiers’ IT.

## Author Contributions

MAA and RDD conceived and designed the experiments; IJT implemented DAStk and performed the experiments; IJT and RDD analyzed the data; all authors contributed to writing the paper.

## Abbreviations

The following abbreviations are used in this manuscript:

TF: Transcription factor
ATAC: Assay for Transposase-Accessible Chromatin
SNP: single nucleotide polymorphisms
ChIP: chromatin immunoprecipitation
eRNA: enhancer RNA
MD-score: motif displacement score
SCLC: small cell lung carcinoma

